# Plant Development Drives Dynamic Shifts in the Root Compartment Microbiomes of Wild and Domesticated Finger Millet Cultivars

**DOI:** 10.1101/2024.04.07.588467

**Authors:** Fantaye Ayele Dadi, Saraladevi Muthusamy, Samrat Ghosh, Diriba Muleta, Kassahun Tesfaye, Fassil Assefa, Jie Xu, Farideh Ghadamgahi, Rodomiro Ortiz, Ramesh Raju Vetukuri

## Abstract

**Background:** Plant-microbe interactions in two root compartments - the rhizosphere and endosphere - play vital roles in maintaining plant health and ecosystem dynamics. The microbial communities in these niches are shaped in complex ways by factors including the plant’s developmental stage and cultivar, and the compartment where the interactions occur. Different plant cultivars provide distinct nutritional and ecological niches and may selectively enrich specific microbial populations through the secretion of root exudates. This gives rise to complex and dynamic plant-microbe interactions; some cultivars promote the recruitment of beneficial symbionts while others may deter pathogens. To clarify these processes, this work investigated the structure of the endosphere and rhizosphere microbial communities of wild type finger millet and five domesticated cultivars across two plant developmental stages.

**Results:** Our results showed that the plant developmental stage, compartment, and cultivar have varying degrees of impact on root-associated microbiomes. The dominant bacterial phyla in all samples were *Proteobacteria*, *Actinobacteria*, and *Bacteroidetes*, while the dominant fungal phyla were *Ascomycota* and *Basidiomycota*. All of these phyla exhibited pronounced variations in abundance. In general, an increased abundance of *Actinobacteria* in the endosphere was accompanied by a reduced abundance of *Proteobacteria*. The most pronounced changes in microbial community structure were observed in the rhizosphere during the flowering stage. Changes in the microbiome patterns of the rhizosphere were driven predominantly by the genus *Pseudomonas.* Moreover, the host plant’s developmental stage strongly influenced the microbial communities, suggesting that plants can recruit specific taxa based on their need for particular soil consortia.

**Conclusions:** Our results show that both host developmental stage and domestication strongly affect the assembly and structure of the plant microbiome. Moreover, plant root compartments can selectively recruit specific taxa from associated core microbial communities to fulfill their needs in a manner that depends on both the plant’s developmental stage and the specific root compartment that is involved. These findings show that deterministic selection pressures exerted by plants during their growth and development can significantly affect their microbial communities and have important implications for efforts to create tools for manipulating the microbiome to sustainably improve primary productivity.

## Background

Finger millet (*Eleusine coracana*) is an essential cereal crop renowned for its resilience and nutritional value, particularly in semiarid and tropical regions. It is the sixth most important cereal crop globally, with annual production of around 4.5 million metric tons, and has been cultivated since the establishment of the earliest indigenous African communities [1]. Predominantly cultivated in African and Asian countries, it serves as a vital staple source of food and feed for populations in developing nations [2–4]. Millet grains are rich in essential macronutrients, minerals, and polyphenols, and their nutritional value exceeds that of conventional cereals such as rice and wheat [3, 5]. Moreover, the resilience of finger millet in harsh environments with poor soil fertility and elevated salinity allows it to flourish in areas where conventional agriculture is impractical.

This adaptability of finger millet plants is related to their interactions with soil microorganisms in plant compartments within the roots, stems, and seeds. These interactions are influenced by the unique microenvironment of each compartment [1]. The microbiota of the root often (but not always) contains a greater diversity of microorganisms than those of other plant compartments [2]. The rhizosphere and endosphere are the most prominent root compartments [3]; the rhizosphere comprises the root surface and adjacent soil particles, while the endosphere comprises the roots’ internal tissue [4]. They have different environmental conditions resulting from differences in their carbon sources, content of host plant-derived compounds, and exposure to climatic factors [5, 6]. Consequently, the composition of the rhizosphere microbiota differs significantly from that of the endosphere.

The bacteriota (a component of the microbiota) of the rhizosphere is the consortium of bacteria residing in the soil around the roots and on the root surfaces. Its development is significantly influenced by a complex process of rhizodeposition [7–10] during which a subset of microorganisms from the rhizosphere infiltrates plant roots and colonizes the endosphere. This process is partly controlled by the plant’s intrinsic immune system and causes the two rhizocompartments to harbor unique microbial communities with differing compositions and structures that are partly controlled by host-driven mechanisms [11]. The plant roots host diverse microorganisms that may include both plant growth-promoting (beneficial) microbes and potentially harmful pathogens that can be neutralized through interactions with host plants. Some beneficial bacterial species have been shown to actively enhance plant growth, health, and development by facilitating nutrient acquisition, performing hormone synthesis, mitigating various stresses, and protecting against pathogen infection [7, 11]. For example, one study indicated that Phl-producing pseudomonads isolated from the finger millet rhizosphere could have applications in the biocontrol of phytopathogens [12]. Another study showed that incorporating arbuscular mycorrhizal fungi (AMF) and plant growth-promoting rhizobacteria (PGPR) into the rhizosphere improved crop performance and agricultural sustainability, particularly under stressful conditions [13].

Plant-associated microbiomes substantially influence plants’ phenotypic adaptability and the functionality of their root systems [14, 15]. The composition of these microbial communities may depend on several factors including the soil type [16], environmental factors [17, 18], the cultivar under consideration [19], and the composition of the root exudates, which may include hormones and primary metabolites (e.g., amino acids, organic acids, and sugars) [20, 21]. In general, the microbial activity and biomass content of the rhizosphere microbiome exceed those of the bulk soil [8, 9]. However, for some crop species the microbial activity in the rhizosphere may be similar to that in the soil [22].

Several recent studies [21, 23–25] on the microbiomes of the rhizocompartments (i.e., the endosphere and rhizosphere) and soil in various crops have provided valuable insights into the structure and function of the microbial communities in and around the roots. The bacterial phyla most commonly found in the rhizosphere are *Proteobacteria*, *Acidobacteria*, *Verrucomicrobia*, *Bacteroidetes*, *Planctomycetes*, and *Actinobacteria* [26], while *Actinobacteria* are more dominant in the endosphere [27] along with the fungal phyla *Ascomycota* and *Basidiomycota* [27]. In some cases, a subpopulation of the rhizosphere enters the roots and joins the endosphere microbiome. The composition of the microbiota is plant-specific and may change as the plant progresses through different developmental stages. There is also clear evidence that genetic and cultivar-specific factors can have strong effects on microbial communities and their structure in diverse crops [17, 28]. Similarly, the soil environment and biotic or abiotic stresses can drive shifts in the composition of the root-associated microbiota [1, 20], and microbial interactions can influence the composition of the root endophytic microbiota [29]. Because the composition of the microbiota and its relationship with the host plant are complex, dynamic, and plant-specific, we aimed to determine how the host plant’s developmental stage and genetic background shape the root-associated microbiota in finger millet. To this end we screened the microbiota of wild type finger millet and five domesticated cultivars, focusing on the bacteriota and mycobiota and their changes during the seedling and flowering growth stages.

## Materials and methods

### Finger millet sources and growth conditions

African finger millet plant species, including the wild type *Eleusine coracana* subsp. *Africana* (referred to as Africana in the following text and figures) and domesticated *Eleusine coracana* subsp. *Coracana* (represented by the cultivars Axum, Wama, Padet, Tesema, and Tadesse) were obtained from the Melkassa Research Center, Ethiopian Institute of Agricultural Research (EIAR), Ethiopia. Detailed information on these varieties is given in Table 1. Finger millet seeds were subjected to surface disinfection before sowing to eliminate microbes not present within the endophytic seed-transmittable microbiome. Plants were cultivated without fertilizer in 7.5-liter pots with three plants per pot under controlled conditions in the SLU Biotron chamber in Alnarp, Sweden. The growth environment featured day and night temperatures of 30/27°C, a 12-hour light/dark cycle, and a light intensity of 350 µmol m^-2^ s^-1^. Three biological replicates of plants from each cultivar and treatment were grown.

**Table 1.**
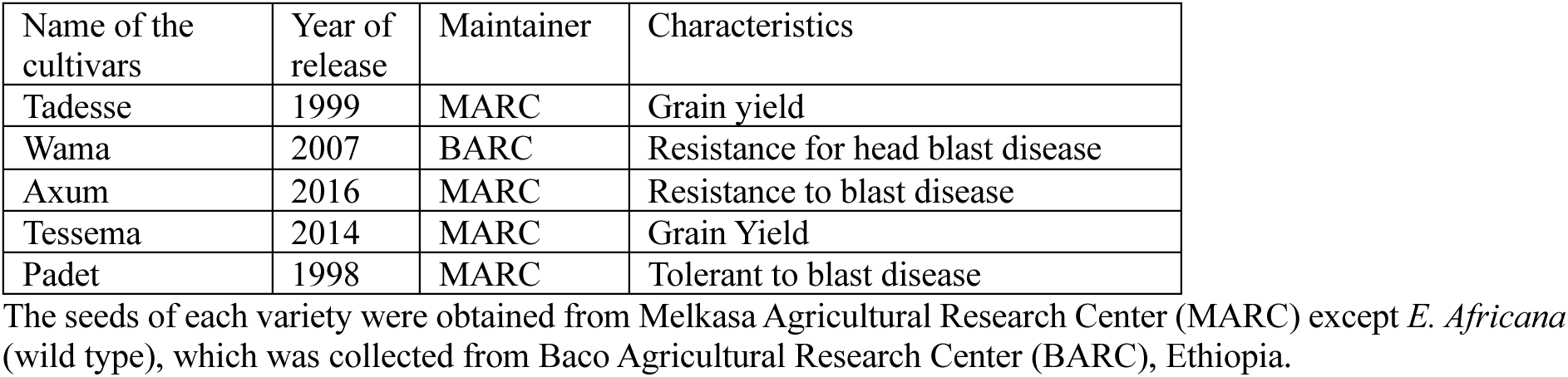
List of bacterial species present only in the five finger millet cultivars during the seedling stage in the endosphere.

| Name of the cultivars | Year of release | Maintainer | Characteristics |
| --- | --- | --- | --- |
| Tadesse | 1999 | MARC | Grain yield |
| Wama | 2007 | BARC | Resistance for head blast disease |
| Axum | 2016 | MARC | Resistance to blast disease |
| Tessema | 2014 | MARC | Grain Yield |
| Padet | 1998 | MARC | Tolerant to blast disease |

The seeds of each variety were obtained from Melkasa Agricultural Research Center (MARC) except *E. Africana* (wild type), which was collected from Baco Agricultural Research Center (BARC), Ethiopia.

## Sample collection

### Collection of root endosphere and rhizosphere samples

Samples from each plant were collected at two different growth stages for metagenomic analysis targeting the 16S rRNA gene and internal transcribed spacer (ITS) region. Rhizosphere and root endosphere samples were collected twice during the growing season: first during the vegetative (seedling) stage, eight weeks after sowing, and second during the flowering stage, five months after sowing, after the plants had shifted from the vegetative to the reproductive stage. The experiment was run from March to July 2022. For sampling, three plants per cultivar were excavated from each pot and the roots were collected. The non-rhizosphere (loosely attached) soil was separated from the roots by shaking. The collected root samples were placed in 50 mL tubes containing 35 mL of sterile phosphate buffer solution (6.33 g/L NaH_2_PO_4_, 8.5 g/L Na_2_HPO_4_ anhydrous, 200 µL/L Tween 20, pH 6.5). The tubes were vigorously shaken for 2 minutes to release rhizosphere soil from the roots, after which the roots were removed, blotted on paper towels, and placed in new, labeled 50 mL tubes for endosphere sampling. The tubes containing rhizosphere soil and excised root samples were placed on dry ice and transported to the laboratory for further processing. Control samples were obtained by collecting 250 g of soil from each pot before sowing for DNA extraction.

## Sample preparation

Root samples were surface sterilized following the method of McPherson et al. 2018 [30]. Initially, the roots were washed with 50% bleach and 0.01% Tween 20 to remove physical impurities and vigorously shaken for 30 seconds. The root segments were then rinsed with 35 mL of 70% ethanol for one minute, rinsed with sterile water three times, and dried on clean paper towels. The effectiveness of the sterilization was confirmed by culturing aliquots from the final wash on potato dextrose agar and nutrient agar (Sigma-Aldrich). Rhizosphere samples were filtered through a sterile 100 µm mesh filter unit (Fisher Scientific, Waltham, MA, USA) into a sterile 50 mL tube, pelleted at 3000× *g* for 10 minutes at room temperature, and then resuspended in phosphate buffer. The rhizosphere was pelleted again by centrifuging the tubes at 15000× *g* at 4°C for 10 minutes. Finally, the rhizosphere pellet was stored at -20°C until DNA extraction.

### DNA extraction and next-generation sequencing (NGS)

DNA was extracted from 36 endosphere and 36 rhizosphere soil samples using the DNeasy Power Soil Pro Kit (Qiagen, CA, USA) following the manufacturer’s instructions. Root endosphere samples were briefly ground to powder using a Mixer Mill MM 400 Shaker (Retsch, UNSPSC, UK) for 5 minutes with glass beads in a grinding jar. Lysis buffer C1 was then added to the pelleted rhizosphere soil and ground root samples, which were subsequently homogenized with FastPrep-24 (MP Biomedicals, USA). The quality and quantity of the extracted DNA was assessed using a Nanodrop (Nanodrop 8000 Thermo Fisher Scientific, USA). For bacteria, the V5-V6 regions of the bacterial 16S rRNA gene were amplified using the primer pair 799F (AACMGGATTAGATACCCKG) and 1115R (AGGGTTGCGCTCGTTG). For fungi, the internal transcribed spacer 2 (ITS2) region was targeted by the primer pair fITS7 (GTGARTCATCGAATCTTTG) and ITS4 (TCCTCCGCTTATTGATATGC) [31, 32]. For each sample, PCR reactions were performed using a 20 μL mixture containing 1x MyTaq buffer containing 1.5 units of MyTaq DNA polymerase (Bioline GmbH, Luckenwalde, Germany), 2 μl of BioStabII PCR Enhancer (Sigma‒Aldrich Co.), ∼1-10 ng of template DNA, and 15 pmol of the appropriate forward and reverse primers. PCR was performed using the following program: predenaturation at 96°C for 1 minute, denaturation at 96°C for 15 seconds, annealing at 55°C for 30 seconds, and extension at 70°C for 90 seconds, followed by 30-33 cycles of amplification for prokaryotic genes and 35-40 cycles for eukaryotic genes. The amplified PCR products were purified with Agencourt AMPure beads (Beckman Coulter, Brea, CA, United States). Illumina libraries were constructed using the Ovation Rapid DR Multiplex System 1-96 (NuGEN Technologies, Inc., California, USA) with 100 ng of purified amplicon pool DNA for each type. After library construction, the Illumina libraries (Illumina, Inc., CA, USA) were combined and subjected to size selection by preparative gel electrophoresis. Sequencing was performed on an Illumina MiSeq platform at the LGC’s sequencing facility in Berlin (LGC Genomics GmbH, Germany).

### Bioinformatics and statistical analysis

Sequence data analysis was performed using the open-source bioinformatics tool QIIME 2 [33]. Briefly, adapter and primer sequences were trimmed off using the cutadapt plugin [34] and the trimmed sequences were processed with the dada2 plugin [35]. All identified amplicon sequence variants (ASVs) were analyzed further. To classify bacteria, a naïve Bayes classifier was trained using the V5-V6 region of the reference sequences from Greengenes 13_8 (99% sequence similarity) using the QIIME2 plugin feature classifier [36]. Fungal classification was performed using a pre-trained classifier on the UNITE reference database provided by the QIIME2 development team.

Alpha diversity was estimated with Shannon’s diversity index, beta diversity was calculated using the Bray‒Curtis distance matrix, and PERMANOVA was used for statistical testing. Differential gene abundance analysis was performed using LEfSe [37]. Core microbiome analysis and network analysis were performed using the UpSetR [38] and SpiecEasi R [39] packages, respectively. P-values were adjusted for multiple comparisons using the Benjamini–Hochberg method with a statistical significance threshold of P < 0.05.

## Results

### Microbial composition of the endosphere and rhizosphere during the seedling and flowering stages

In all cultivars and control soil samples, the dominant bacterial phyla were *Proteobacteria*, *Bacteroidetes*, and *Actinobacteria* (Fig. 1A, Additional file1). However, *Proteobacteria* was more abundant in the rhizocompartments than in the soil, while the opposite was true for *Bacteroidetes*. There were also notable differences in the relative abundance of the dominant phyla between plant compartments and growth stages. For example, *Actinobacteria* were more abundant in the endosphere than the rhizosphere. In addition, there was a clear shift in the composition of the microbiota of the rhizosphere between the seedling and flowering growth stages, and there were notable compositional differences between the cultivars (Fig. 1A). For example, during the flowering stage the abundance of *Proteobacteria* in the rhizosphere was significantly higher in the five domesticated cultivars than in the wild type (Fig. 1A). Compositional shifts were also observed at lower taxonomic levels, as shown in Fig. 1C. For example, *Chitinophaga* (phylum *Bacteriodetes*) was among the ten most abundant genera in all samples but was substantially more abundant in the control soil than in the root-associated bacteriota. Additionally, the *Pseudomonas* genus exhibited low abundance during the seedling stage in both root compartments (Fig. 1C, Additional file1) but its abundance in the rhizosphere increased dramatically in the flowering stage. Genotype-specific variations were also observed. For example, during the seedling stage, high abundances of unidentified genera belonging to *Xanthomonadaceae* were detected in the endosphere of cv. Tessema and Tadesse. During the seedling stage, high levels of these genera were also detected in the rhizosphere of the two cultivars mentioned above as well as the Padet and Axum cultivars. In addition, an unidentified genus belonging to *Enterobacteriaceae* exhibited high abundance in the rhizosphere of cv. Tessema and Tadesse during the flowering stage.

**Figure 1.**
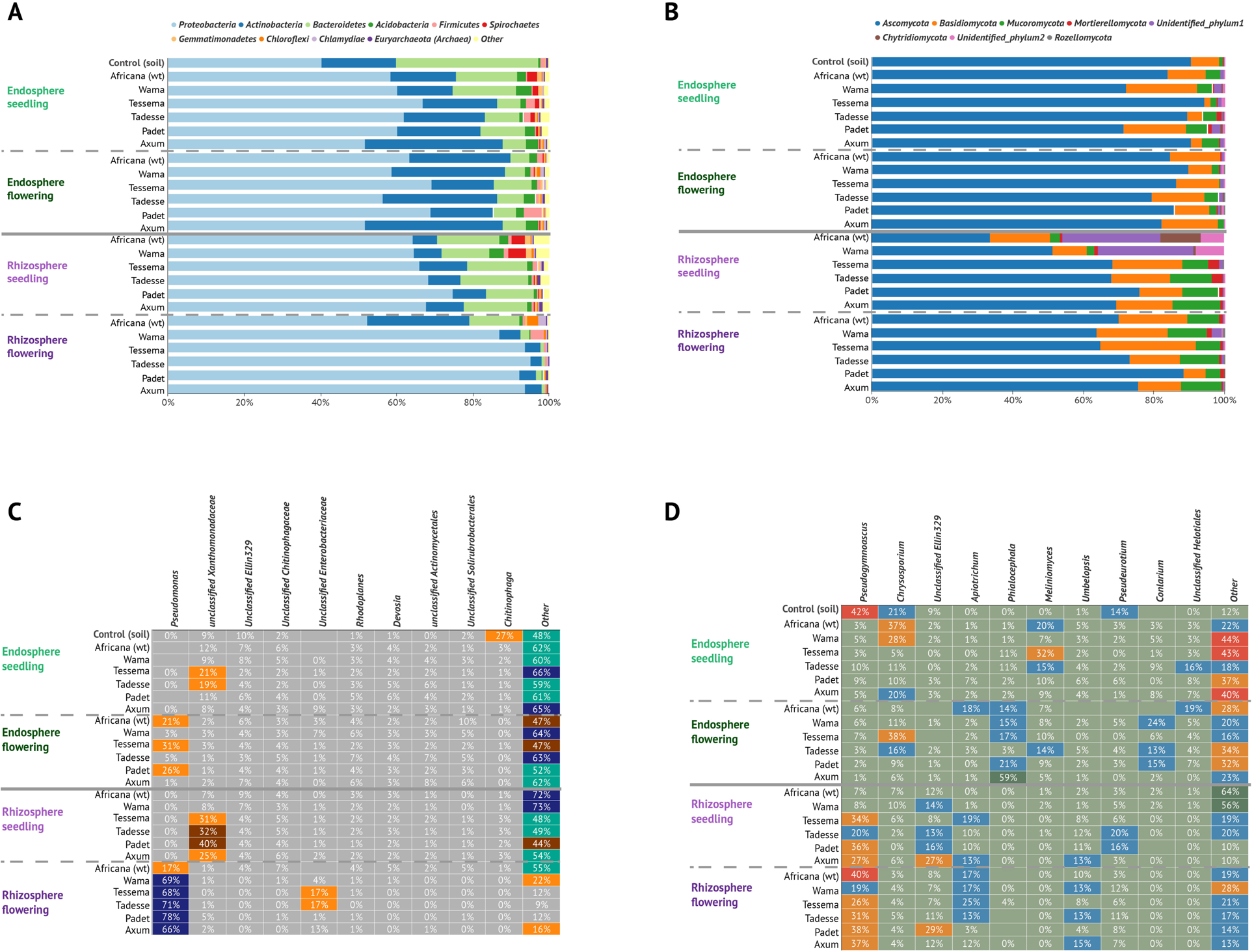
Comparative Analysis of Bacteriota and Mycobiota Across Plant Compartments. This figure shows the top ten bacterial phyla (**A**), all detected fungal phyla (**B**), the top ten bacterial genera (**C**), and the top ten fungal genera (**D**) identified in the endosphere and rhizosphere of plants during the seedling and flowering stages. The samples compared include control (soil without plantation), wild type (Africana), and five different cultivars (Wama, Tessema, Tadesse, Padet and Axum). Relative abundances > 0 are shown in the heatmaps of (**C**) and (**D**).

In all samples, the most abundant fungal phyla in the mycobiota were *Ascomycota* and *Basidiomycota*, followed by *Mucoromycota* (Fig. 1B, Additional file 3). Additionally, several unidentified phyla were detected in the rhizosphere of the wild type and cv. Wama during the seedling stage. In all growth stages, *Ascomycota* were less abundant in the rhizosphere than in the endosphere.

At the genus level, the soil samples were dominated by genera including *Pseudogymnoasucus*, *Chrysosporium*, and *Pseudeurotium* (Fig. 1D, Additional file 4). Rhizosphere samples were dominated by *Pseudogymnoasucus*, *Apiotrichum*, *Umbelopsis* and *Unclassified Ellin329* (all belonging to the phylum *Ascomycota*) in both plant developmental stages (Fig. 1D). The endosphere samples had a distinct set of dominant genera that included *Chrysosporium*, *Meliniomyces*, *Conlarium*, and *Philaocephala* (Fig. 1D).

### Similarities and differences between cultivars in different plant developmental stages and root compartments

A species-level core microbiome analysis was conducted to identify shared and unique microbial species. In the endosphere seedling stage, 94 bacterial species were shared among all the sample types including the control soil samples (Fig. 2A). Fifty species were detected among the cultivars but not in the control soil, and four were shared exclusively among the cultivars but not in the wild type (Fig. 2A and Table 1). For the endosphere flowering stage samples, 43 bacterial species were commonly detected in all cultivars and the wild type, and five species were detected in all cultivars but not the wild type (Table 2).

**Figure 2.**
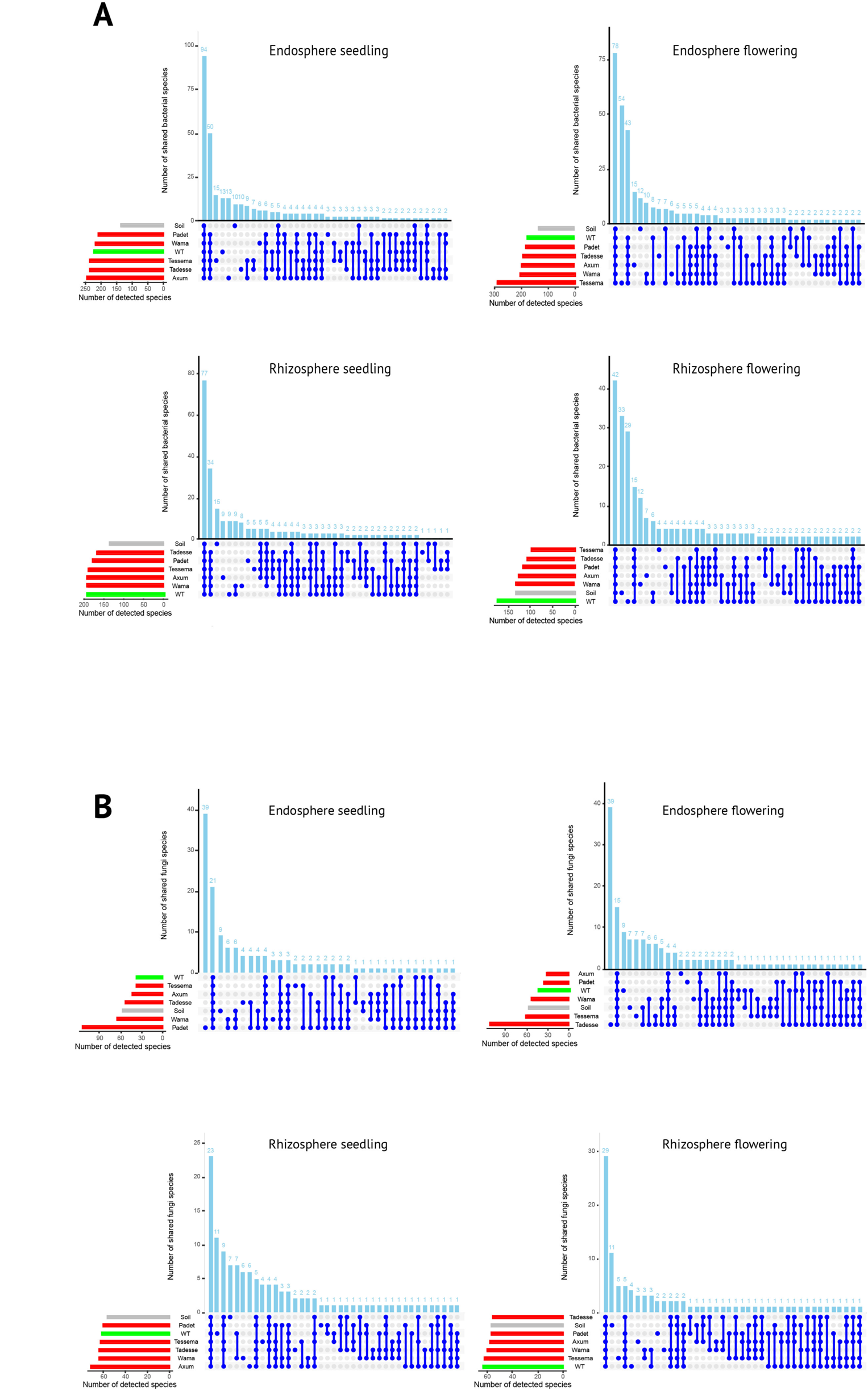
UpSet plot showing the results of core microbiome analyses of the bacterial species (**A**) and fungal species (**B**) present in the endosphere and rhizosphere of wild type plants (WT; Africana), five different cultivars (Wama, Tessema, Tadesse, Padet, and Axum), and control soil (soil without plantation) during the seedling and flowering growth stages. The number of detected and shared species are indicated by the bars in the lower left corner and the columns at the top, respectively. Connected dots represent species shared among sample types (control, wild type, and cultivars), while single dots represent species unique to a specific sample type.

**Table 2.** List of bacterial species present only in the five finger millet cultivars during the flowering stage in the endosphere.

|  | Phylum | Class | Order | Family | Genus | Species |
| --- | --- | --- | --- | --- | --- | --- |
| 1 | *Proteobacteria* | *Gammaproteobacteria* | *Enterobacteriales* | *Enterobacteriaceae* |  |  |
| 2 | *Proteobacteria* | *Betaproteobacteria* | *Burkholderiales* | *Burkholderiaceae* | *Burkholderia* |  |
| 3 | *Proteobacteria* | *Betaproteobacteria* | *Burkholderiales* | *Oxalobacteraceae* | *Telluria* | *mixta* |
| 4 | *Proteobacteria* | *Gammaproteobacteria* | *Oceanospirillales* | *Halomonadaceae* | *Halomonas* | *phosphatis* |

In the rhizosphere seedling samples, 34 bacterial species were found in all finger millet samples (including the wild type) and two were found only in the cultivars (Table 3). The wild type had a distinctive microbial community composition in the rhizosphere during the seedling stage, with nine unique species (Fig. 2A). A shift in the core rhizosphere microbiome was observed in the flowering stage: 20 unique bacterial species were found in the wild type, and 15 were found in all finger millet genotypes but not in the control soil. Moreover, three species were found in all of the cultivars but not in the wild type or control samples (Table 4).

**Table 3.** List of bacterial species present only in the five finger millet cultivars during the seedling stage in the rhizosphere.

|  | Phylum | Class | Order | Family | Genus | Species |
| --- | --- | --- | --- | --- | --- | --- |
| 1 | *Proteobacteria* | *Deltaproteobacteria* |  |  |  |  |
| 2 | *Chloroflexi* | *TK10* | *TK10* |  |  |  |
| 3 | *Gemmatimonadetes* | *Gemmatimonadetes* | *Gemmatimonadetes* |  |  |  |
| 4 | *Proteobacteria* | *Alphaproteobacteria* | *Rhodobacterales* | *Rhodobacteraceae* | *Rhodobacter* |  |
| 5 | *Proteobacteria* | *Actinobacteria* | *Actinomycetales* | *Nocardiaceae* | *Rhodococcus* |  |

**Table 4.** List of bacterial species present only in the five finger millet cultivars during the flowering stage in the rhizosphere.

|  | Phylum | Class | Order | Family | Genus | Species |
| --- | --- | --- | --- | --- | --- | --- |
| 1 | *Proteobacteria* | *Gammaproteobacteria* | *Legionellales* | *Legionellaceae* |  |  |
| 2 | *Proteobacteria* | *Alphaproteobacteria* | *Rhizobiales* | *Methylocystaceae* |  |  |

In general, fewer fungal species were shared among the studied samples than bacterial species (Fig. 2B). In the endosphere seedling stage, the Padet cultivar had more unique species (39) than the other cultivars. Twenty-one species were found in all sample types including the control soil samples (Fig. 2B), while four fungal species were found in all finger millet samples (including the wild type) but not the control soil. No commonly detected species were present in all five cultivars (Fig. 2B). In the endosphere flowering stage, the Tadesse cultivar had a greater number of unique species than the other cultivars, and only two species were commonly present in all cultivars during the flowering stage (Table 5). In the rhizosphere seedling stage, four fungal species were common to all cultivars (Table 6) and the wild type had the highest number of unique fungal species (Fig. 2B). In the rhizosphere flowering stage, five fungal species were present in all cultivars and the wild type, and only a single species was commonly present in the five cultivars but not the wild type (Table 7) (Fig. 2A).

**Table 5.** List of fungal species present only in the five finger millet cultivars during the flowering stage in the endosphere.

|  | Phylum | Class | Order | Family | Genus | Species |
| --- | --- | --- | --- | --- | --- | --- |
| 1 | *Proteobacteria* | *Gammaproteobacteria* | *Enterobacteriales* | *Enterobacteriaceae* |  |  |
| 2 | *Proteobacteria* | *Gammaproteobacteria* | *Pseudomonadales* | *Pseudomonadaceae* | *Pseudomonas* |  |
| 3 | *Proteobacteria* | *Alphaproteobacteria* | *Rhizobiales* | *Rhizobiaceae* | *Rhizobium* |  |

**Table 6.**
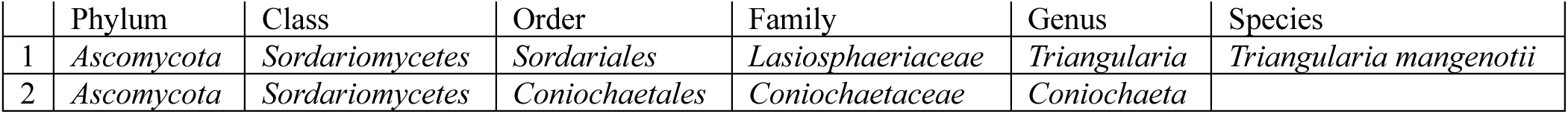
List of fungal species present only in the five finger millet cultivars during the seedling stage in the rhizosphere.

**Table 7.** The fungal species present only in the five finger millet cultivars during the flowering stage in the rhizosphere.

|  | Phylum | Class | Order | Family | Genus | Species |
| --- | --- | --- | --- | --- | --- | --- |
| 1 | *Ascomycota* | *Sordariomycetes* | *Sordariales* | *Cephalothecaceae* | *Phialemonium* | *Phialemonium inflatum* |
| 2 | *Ascomycota* | *Sordariomycetes* | *Sordariales* | *Cephalothecaceae* | *Phialemonium* | *Phialemonium atrogriseum* |
| 3 | *Ascomycota* | *Sordariomycetes* | *Sordariales* | *Lasiosphaeriaceae* | *Triangularia* | *Triangularia mangelotii* |
| 4 | *Ascomycota* | *Eurotiomycetes* | *Eurotiales* | *Trichocomaceae* | *Rasamsonia* | *Rasamsonia byssochlamydoides* |

**Table 8.**

|  | Phylum | Class | Order | Family | Genus | Species |
| --- | --- | --- | --- | --- | --- | --- |
| 1 | *Ascomycota* | *Sordariomycetes* | *Hypocreales* |  | *Acremonium* | *A.alternatum* |

### Enrichment of specific microbial taxa during the seedling and flowering stages

Differential abundance analysis of the bacterial microbiota using linear discriminant analysis effect size (LEfSe) revealed the specific taxa responsible for differences in the composition of the microbiota between growth stages in each rhizocompartment (Fig. 3). In the endosphere, the most pronounced dissimilarities in the bacterial microbiota were seen during the seedling stage, where significant enrichment of multiple taxa was observed in the wild type and three cultivars (Fig. 3A). In the wild type, 26 taxa were enriched and more than 11 had a large effect size (LDA score > 7); the most enriched taxon was the class *Saprospirae* (LDA score > 7). The Wama cultivar had 23 significantly enriched taxa, with the candidate order *Elin329* having the largest effect size (LDA score > 7). In Padet, the *Halomonas*, *Phosphatis* and *Oceanospirillales* taxa were significantly enriched (LDA score > 6). The Axum cultivar had seven enriched taxa, with the genus *Rhodoplanes* having the largest effect size. Conversely, during the flowering stage, only one cultivar exhibited differential enrichment in the endosphere: cv. Padet had four significantly enriched taxa (Fig. 3B).

**Figure 3.**
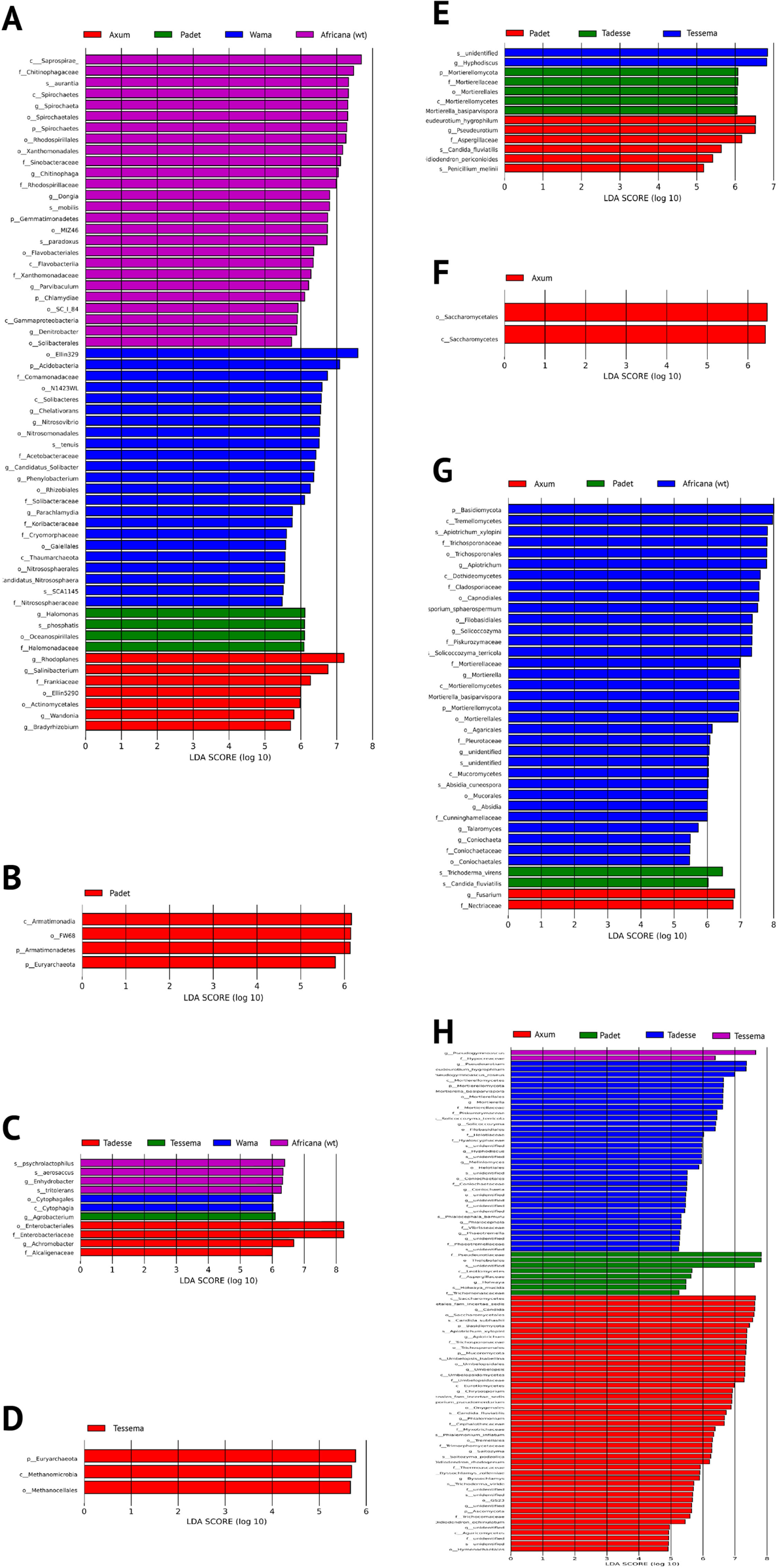
Differential Abundance Analysis Using LEfSe. This figure shows the differentially abundant bacterial taxa (**A-D**) and fungal taxa (**E-H**) identified in wild type finger millet (wt_Africana) and five cultivars (cv. Wama, cv. Tessema, cv. Tadesse, cv. Padet, cv. Axum). Analyses were conducted for each compartment in each growing stage: Endosphere seedling (**A, E**), Endosphere flowering (**B, F**), Rhizosphere seedling (**C, G**), and Rhizosphere flowering (**D, H**). Taxa with linear discriminant analysis (LDA) scores > 4 at p < 0.05 were considered to exhibit significant differential abundance.

In the rhizosphere, the wild type and three cultivars exhibited significant enrichment of bacterial taxa during the flowering stage (Fig. 3C); the wild type and cv. Tadesse had the highest number of enriched taxa, followed by cv. Wama and cv. Tessema (Fig. 3C). Conversely, only cv. Tessema exhibited significant enrichment during seedling stage; *Euryarchaeota*, *Methanomicrobia*, and *Methanocellales* had the largest effect sizes (LDA > 5; Fig. 3D). The most pronounced dissimilarities in the mycobiota were observed in the rhizosphere. Four cultivars had significantly enriched taxa during the seedling stage; cv. Axum (47) and cv. Taddesse (35) had the highest numbers of significantly enriched taxa, followed by cv. Padet and cv. Tessema (Fig. 3H). The wild type had more enriched taxa during the flowering stage than the other cultivars. *Basdiomycota* had the largest effect size, with an LDA score > 8. The Padet and Axum cultivars harbored two enriched taxa each (Fig. 3G). In the endosphere, only the Axum cultivar had significantly enriched taxa during the flowering stage; the largest effect size (LDA > 6) was observed for *Saccharomycetales* members LDA > 6 (Fig. 3F). In addition, three cultivars had significantly enriched taxa during the seedling stage. The *Hyphodiscus* genus had the largest effect size in cv. Tessema (LDA score > 6), while the Tadesse and Padet cultivars both had 4 enriched taxa (Fig. 3E).

### Microbial diversity comparisons between cultivars and plant developmental stages

The endosphere bacteriota had similar levels of alpha diversity across all cultivars and the wild type during both the seedling and flowering stages Fig. 4A. However, cv. Tessema had slightly lower alpha diversity in the endosphere during the flowering stage (Fig. 4A). The Padet cultivar had the lowest alpha diversity in the rhizosphere. Overall, the wild type maintained a high level of alpha diversity in the rhizosphere even during the flowering stage, when the other cultivars had significantly less diversity. However, there were no significant differences between cultivars with respect to the beta diversity of the bacterial and fungal communities for any rhizocompartment or developmental stage.

**Figure 4.**
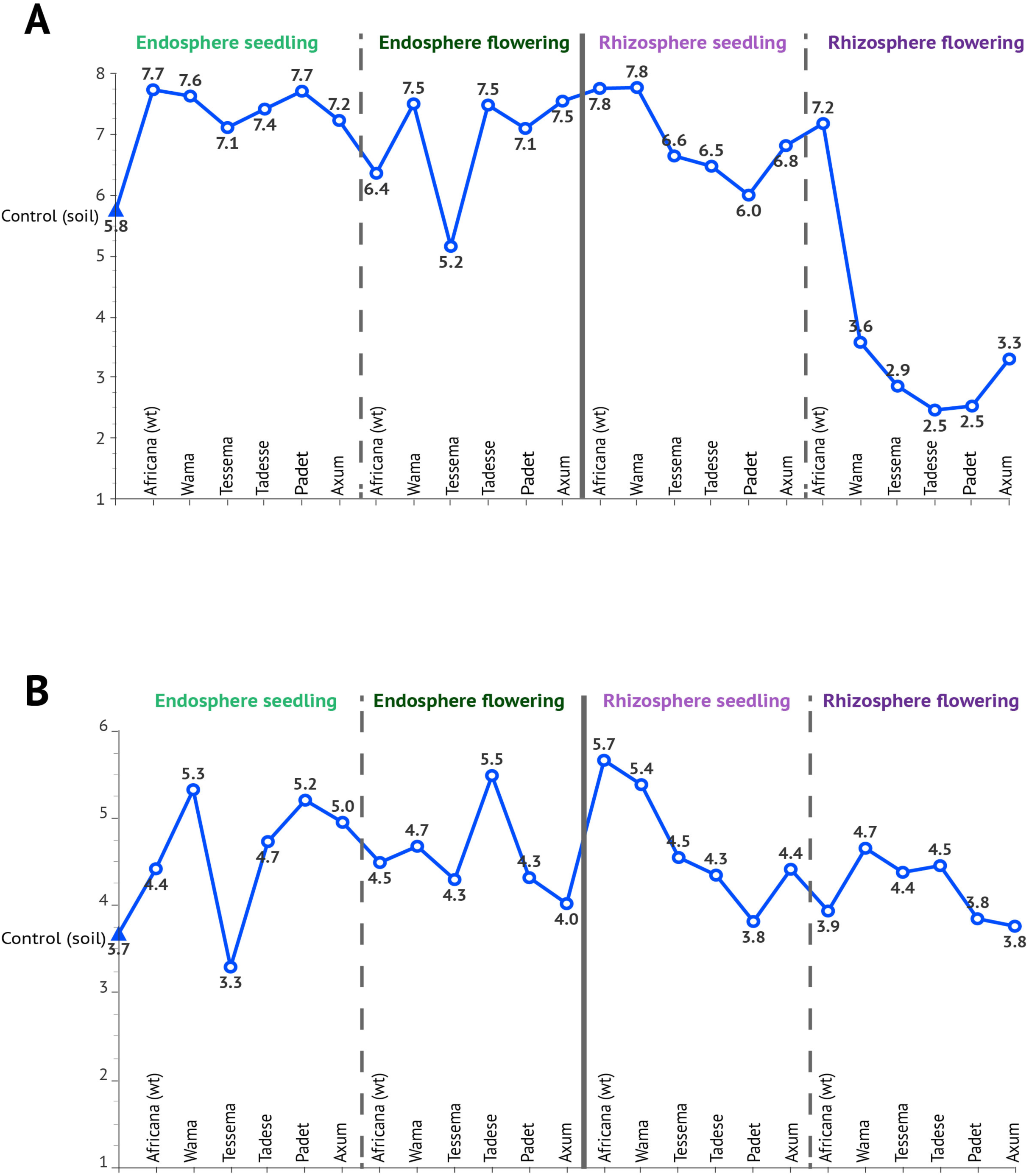
Changes in Alpha Diversity of Bacteriota and Mycobiota. This figure shows the alpha diversity of bacteriota (**A**) and mycobiota (**B**), estimated using Shannon’s diversity index. The alpha diversity in the rhizosphere was clearly lower than in the endosphere during both the seedling (p = 0.049) and flowering stages (p < 0.001). Moreover, the alpha diversity of the rhizosphere declined significantly during the flowering stage (p < 0.001). No significant changes were observed in the endosphere or among with mycobiota (p = 0.186). The data are represented using median values for each sample type including control (soil without plantation), wild type (Africana), and five domesticated cultivars (Wama, Tessema, Tadesse, Padet, and Axum). All p-values were adjusted for multiple comparisons.

The alpha diversity of the rhizosphere and endosphere samples from the finger millet cultivars differed markedly from that in the control soil samples, as shown in Figure 4A. In addition, there were significant differences in the diversity of the bacterial communities in different plant developmental stages and rhizocompartments (Kruskal-Wallis test, p-value <0.001). Specifically, the rhizosphere exhibited lower microbial diversity during the flowering stage than the other compartments (including the endosphere and control soil). However, the diversity of the rhizosphere bacterial community in the seedling stage was higher and comparable to that of the endosphere (Fig.4A). Remarkably, the wild type finger millet maintained a higher alpha diversity than the cultivars in both the seedling and the flowering stages. In particular, the root endosphere consistently displayed higher levels of alpha diversity than any cultivar other than cv. Tessema irrespective of growth stage (Fig.4A).

In contrast, fungal diversity was relatively consistent across both growth stages and plant rhizocompartments, with only a few cultivars showing variations (Fig. 4B). No significant differences were detected between the groups (Kruskal-Wallis test, p = 0.186). However, the fungal alpha diversity in the rhizosphere during the flowering stage was slightly lower than in the bulk soil and the endosphere. In contrast to the bacterial diversity patterns, the fungal alpha diversity of the wild type finger millet plants fluctuated throughout the study (Fig. 4B). Moreover, the drastic decline in alpha diversity seen in the bacterial community of the rhizosphere during the flowering stage was not seen in the corresponding mycobiota (Fig. 4B). A beta diversity analysis based on the Bray‒Curtis distance matrix revealed significant differences between the rhizosphere and endosphere compartments at different stages of plant development (PERMANOVA, p = 0.001, Fig. 5A). Notably, the clustering pattern of the soil samples differed from that of the wild type and the cultivars. Furthermore, the microbial communities associated with the flowering stage had different clustering characteristics to the seedling-stage plants (Fig. 5A). A beta diversity analysis of the fungal community also revealed significant differences (PERMANOVA, p-value = 0.001) between samples in the rhizosphere (Fig. 5B). As previously observed for the bacterial communities, the pattern of the fungal communities in the rhizosphere differed from those in the endosphere (Fig. 5B).

**Figure 5.**
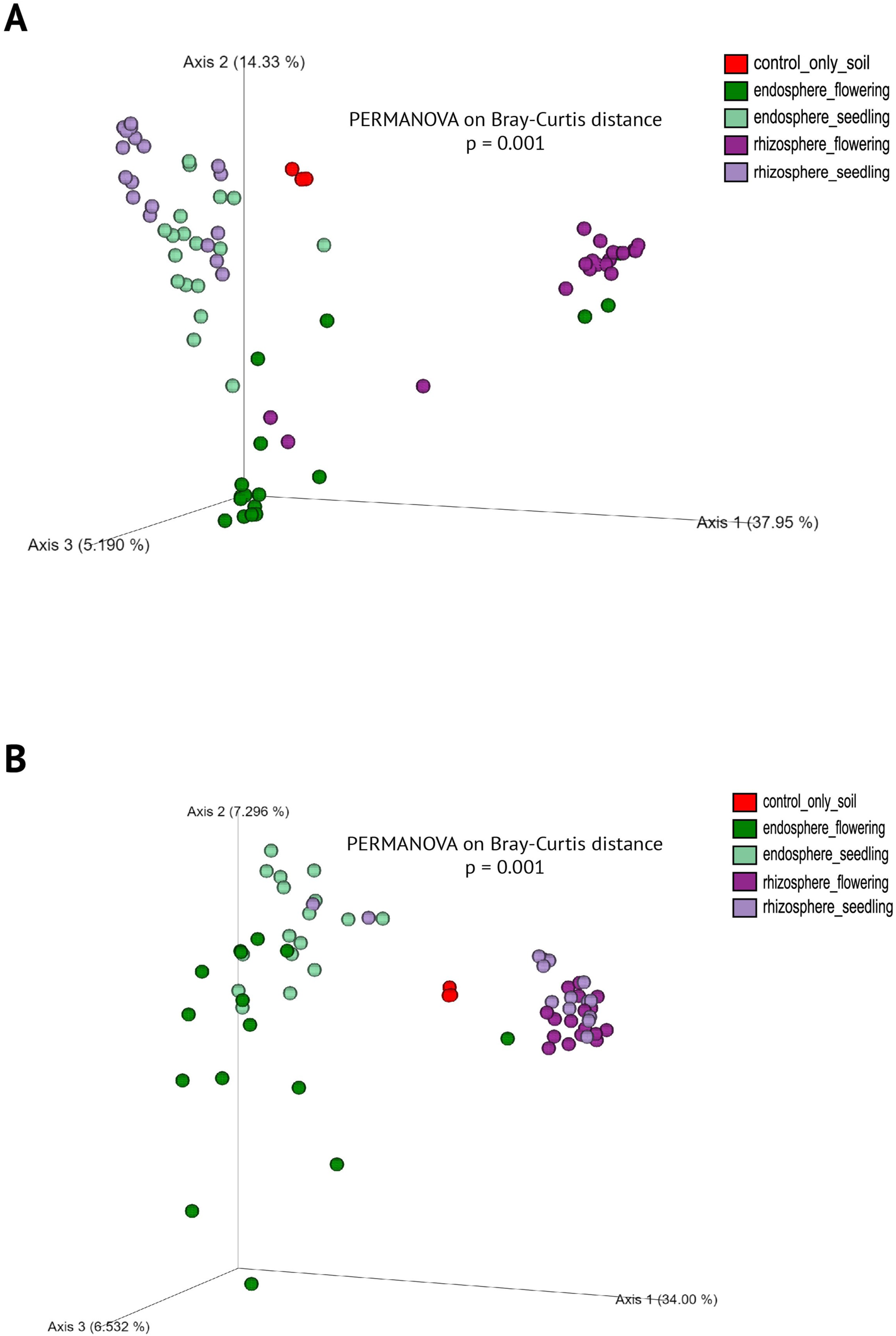
Differences in Beta Diversity of Bacteriota and Mycobiota. This figure shows the beta diversity of bacteriota (**A**) and mycobiota (**B**) in two plant compartments and plant developmental stages. Significant differences were found in all pairwise comparisons (p <0.05; p values were corrected for multiple comparisons).

### Interkingdom species association during plant developmental stages

We conducted a network analysis to detect potential inter-kingdom associations between bacterial and fungal species. Because only a few samples were available for each cultivar, we focused on co-occurrence patterns at the compartment level and across the two plant developmental stages. Fig. 6 shows the network patterns that were detected. In general, intra-kingdom associations are more complex than intra-kingdom associations. However, the flowering stage network in the rhizosphere was very simple (Fig. 6D), with only a single inter-kingdom association – a co-exclude association between nodes 215 (Fungi, *Myxotrichaceae*) and 105 (Bacteria, *Chitinophagceae*). The most complex network was observed in the rhizosphere during the seedling stage (Fig. 6C). The endosphere networks in the seedling and flowering stages were similar and of intermediate complexity (Fig. 6A and 6B). There was also a consolidation of the networks between certain bacterial and fungal species in the endosphere during the flowering stage (Fig. 6B). Detailed taxonomic affiliations of the species and the correlations for their interactions are presented in Additional file A6.

**Figure 6.**
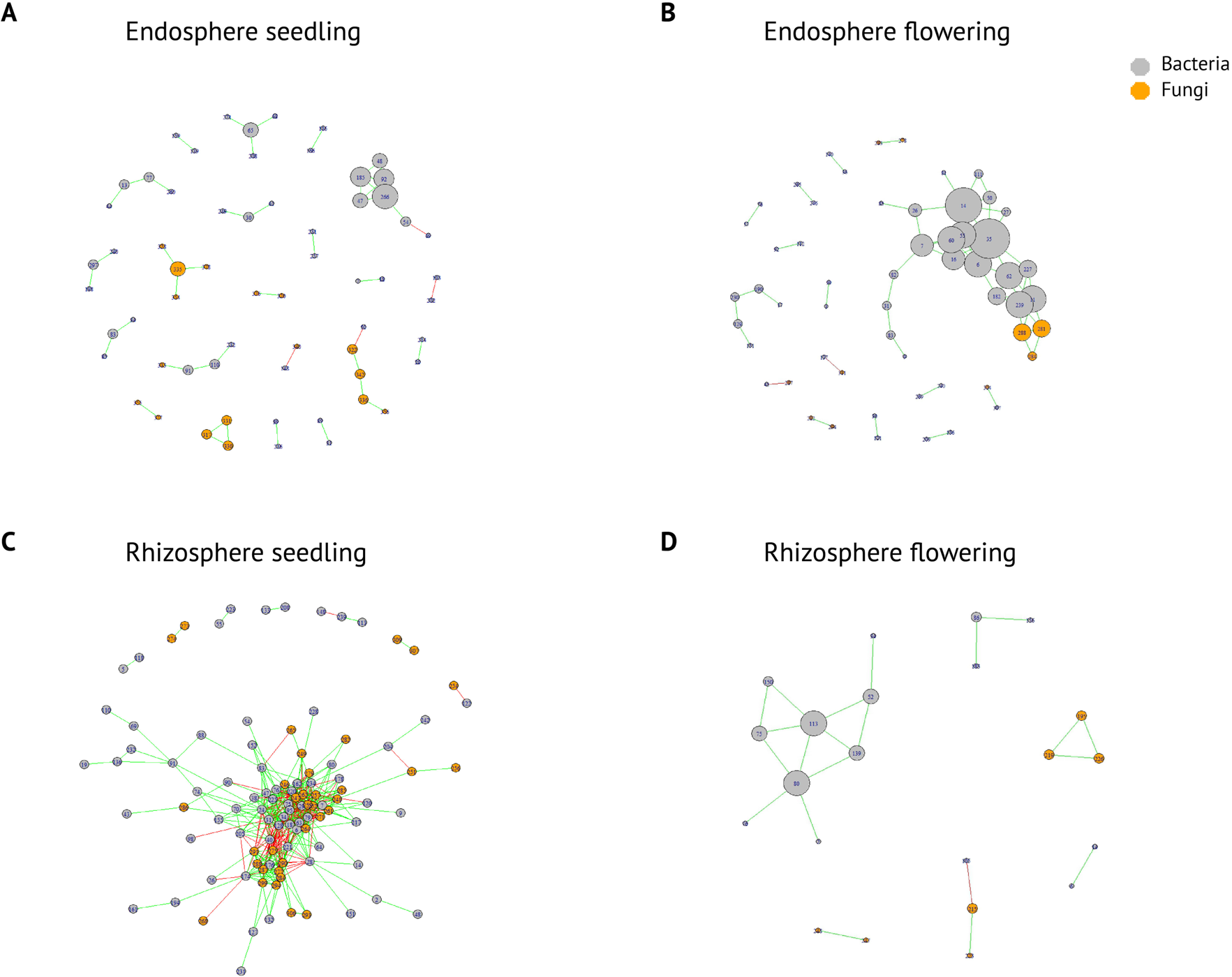
Co-occurrence patterns of bacterial and fungal species during two growth stages in the endosphere and rhizosphere. Only significant associations having |ρ|>0.8 and p<0.05 were plotted. Gray nodes represent bacterial species and orange nodes represent fungal species. Positive correlations are shown in green and negative correlations are shown in red. The sizes of the nodes represent their number of degrees; the bigger the circle, the higher the centrality. Taxonomic information for each node is presented in Table A6.

## Discussion

The complex interplay between soil microbiomes and plant-root-associated microbial communities is crucial in both ecological and agricultural contexts. Research has shown that environmental factors such as the soil type and its physiochemical properties, plant species [15] [19], developmental stage [40], root exudates [20], and domestication [41] intricately shape these microbiomes, affecting plant growth and health [22]. Studies on the microbiota composition and diversity in the endosphere and rhizosphere compartments of olive trees [42], desert plants [43], kiwi fruits [27], and Populus trees [44] have revealed the presence of distinctive root-associated microbiomes but have not characterized their variation over time. In this work, we sought to determine 1) whether the root-associated microbiomes of the rhizosphere and endosphere remain stable over time or undergo compositional changes between the seedling and flowering stages; and 2) whether the root microbiomes of domesticated finger millet cultivars differ from those of the wild type.

Our results show that the structure of the root-associated microbiota in finger millet changes between growth stages and differs significantly between the endosphere and rhizosphere. Moreover, the composition of the root microbiota in domesticated cultivars differs from that of the wild type. These differences in microbial composition were apparent in both the bacteriota and the mycobiota (Fig. 1). For example, the bacterial phyla *Proteobacteria*, *Actinobacteria*, and *Bacteroidetes* dominated the rhizosphere, while *Actinobacteria* were more abundant in the endosphere. *Ascomycota* and *Basidiomycota* were the dominant fungal phyla in both compartments [25, 27, 42, 43]. At a lower taxonomic level, the bacterial genus *Pseudomonas* was enriched in the rhizosphere during the flowering stage and to a lesser degree in the endosphere, while the fungal genus *Pseudogymnoascus* was more abundant in the rhizosphere. These observations show that the microbial communities of the rhizosphere and root endosphere are both dynamic and structurally distinct, as reported previously [45, 46].

Earlier studies have shown that plant microbial communities undergo shifts as the host plant transitions between developmental stages [40, 47]. The composition of the root exudates also changes as the plant advances through its developmental stages, which may be linked to shifts in the composition of the microbial communities [40]. In later growth stages, plants release specific substrates that may encourage specific microbial taxa to enter the rhizosphere [40, 48]. This might occur because certain microbes can help the plant boost its pathogen resistance, innate immune responses, and ability to cope with abiotic stresses [49]. Our data also revealed cultivar-specific differences in microbiota composition between root compartments. For example, during the flowering stage *Pseudomonas* was less enriched in the rhizosphere of the wild type than in the five domesticated cultivars. However, its abundance in the endosphere during the same stage varied markedly among the crops (Fig.1C), being 21% in the wild type, 31% in Tessema, 26% in Padet, 3% in Wama, 5% in Tadesse, and 1% in Axum.

Members of the *Pseudomonas* genus have been found in the rhizospheres of several plants, including finger millet [43, 44, 48]. The efficiency of *Pseudomonas* spp. in controlling pathogens and promoting plant growth is closely related to their ability to competitively colonize the rhizosphere and persist in this compartment – a quality known as rhizocompetence [50]. *Pseudomonas* species are known for their diverse beneficial effects on plants, which include protecting against phytopathogens [12, 50, 51], solubilizing minerals (e.g., phosphate), and promoting plant growth under abiotic stress conditions [51]. For example, this genus can control the blast disease fungus *Pyricularia grisea* by producing antifungal metabolites, siderophores, and hydrolytic enzymes [52]. The *Pseudomonas* genus also has the remarkable ability to survive in diverse environments, including the rhizosphere where plants release rhizodeposits including root exudates containing diverse nutrient-rich substrates such as organic acids, carbohydrates, fatty acids, amino acids, and proteins that support the rhizosphere microbiome [53]. In a study on wheat, the proliferation of *Pseudomonas spp*. was shown to alter rhizosphere diversity under nutrient-limiting conditions [54].

During the seedling stage, an unidentified *Xanthomonadaceae* species was more abundant in the rhizospheres of all cultivars aside from cv. Wama than in the wild type. Two of the cultivars (Tessema and Tadesse) also had higher levels of this species in the endosphere during the flowering stage. It has been suggested that bacteria within the family *Xanthomonadaceae* could be used as biocontrol agents [55, 56]. However, some *Xanthomonas* species are known plant pathogens [57]. We only detected *Xanthomonas* in 4 out of 75 samples, with an average abundance of < 0.06%. Tessema and Tadesse selectively boosted the proliferation of unidentified *Enterobacteriaceae*; some genera from this family, such as *Enterobacter* and *Klebsiella*, are known to promote plant growth through hormone production, phosphate solubilization, nitrogen fixation, and antimicrobial compound production [58, 59]. The relative abundance of the most abundant fungal genera depended on the host plant’s developmental stage and crop genotype (cultivar). Previous studies identified *Pseudogymnoascus* as the most abundant fungal genus in the rhizosphere and showed that its members contribute to nutrient cycling, plant growth promotion, stress tolerance [60, 61], and cellulose degradation. Saprophytic fungi are known to benefit plant growth by enhancing apocarotenoid biosynthesis and stress tolerance. They comprise roughly 39% of the total fungal community in the rhizosphere of turfgrass but are less abundant in the root endosphere [60], which aligns with our observations. In addition, a large proportion of the sequences assigned to the candidate order *Ellin329*, *Apiotrichum,* and *Umbelopsis* were more abundant in the rhizosphere, albeit with appreciable variation between genotypes.

*Chrysosporium*, *Phialocephala, Meliniomyces*, *Conlarium*, and unclassified *Helotiales* taxa exhibited selective enrichment in the endosphere of the studied millet varieties. There is emerging evidence that these endophytic fungi have plant growth-promoting functions [62–65]. The flowering stage in plants is a reproductive stage in which the plant is particularly sensitive to biotic and abiotic environmental stresses, which can significantly impact the plant’s development and yield. The strong enrichment of *Pseudomonas* genus during the flowering stage (relative to the seedling stage) indicates that finger millet recruits specific genera during flowering to overcome stresses and reinforce the plant’s innate immune responses. Identifying the recruited taxa at the species level was unfortunately impossible in this work but could provide valuable insights into the functional and ecological relationships underpinning these effects. Moreover, the dynamic successional shifts observed here in the bacterial and fungal communities during the flowering stage indicate that plants customize their own functional communities as they mature.

A beta-diversity analysis revealed significant differences between the seedling and flowering growth stages but not between the various studied cultivars (Fig. 5), indicating that the growth stage had a greater impact on the structure of the microbiota in the rhizocompartments than the plants’ genetic background. This conclusion was supported by the results of an alpha diversity analysis. For the bacteriota, the alpha diversity in the rhizocompartments was consistently higher than in the bulk soil except in the case of the rhizospheres of the five cultivars during the flowering stage. This dramatic loss of diversity during the flowering stage suggests that strong selective pressure was exerted during domestication. We therefore speculate that these changes in the composition of the bacteriota may be linked to the selected features of the cultivars. It was previously reported that rhizosphere microbial communities generally exhibit greater diversity than those of the endosphere [43]. This is consistent with the dynamic nature of the rhizosphere, which serves as a primary locus of nutrients and energy resources for neighboring microorganisms and thus creates a competitive environment that fosters a highly active microbial community.

In contrast to the earlier report cited above, our results indicated that the bacterial diversity in the endosphere was generally higher than in the rhizosphere, particularly during the flowering stage. This apparent contradiction may again be linked to shifts in microbial community structure driven by growth stage transitions. Such shifts may occur in response to selective processes within the plant compartments and/or changes in nutrient availability, symbiotic relationships, stress, the protection afforded by the endosphere, and other dynamic changes during plant development. Interestingly, the reduced diversity during the flowering stage was accompanied by strong enrichment of a single genus, supporting the hypothesis of intense selection pressure exerted by the host plant.

Analysis of the rhizocompartments’ mycobiota revealed lower overall diversity than for bacteria (Fig. 1). This may be partly because many fungal sequences could not be assigned to known phyla. Nevertheless, significant differences between cultivars were observed, indicating cultivar-specific recruitment of fungal species during plant development. As previously observed for bacteria, the fungal diversity in the rhizosphere was lower than in the endosphere during flowering, although this trend was not statistically significant for fungi. There was also no clear difference in fungal diversity between the wild type and the cultivars (Fig. 1B).

The observation of the distinct microbiota in the rhizocompartments led us to explore potential correlations between bacterial and fungal species. Network analysis revealed that the complexity of microbial interactions in the rhizosphere decreased from the seedling to the flowering stage. This reduction in complexity aligns with the shift in the plant’s focus from rapid growth and development during the seedling stage to reproductive growth during flowering. In general, the intra-kingdom correlations were more common than inter-kingdom correlations. All of the endosphere samples exhibited similar network complexity, suggesting that the inner-tissue environment undergoes less drastic changes than the rhizosphere, where the selection pressure exerted by the root exudates varies strongly between growth stages. Further study is needed to determine whether the observed co-occurrence and co-exclusion patterns are due to nutrient competition alone or linked to finger millet development.

We also investigated cultivar type-associated microbiota changes. Several studies have concluded that plant domestication influences the composition of both the rhizosphere and endosphere microbiomes. For example, a study of the foxtail millet endosphere revealed clear differences in community composition between cultivars [66]. Earlier studies focused heavily on the influence of domestication on rhizosphere compartments and identified no significant differences in microbiome diversity between wild and domesticated plants [6, 24]. However, none of those earlier studies examined the influence of plant domestication on microbial communities in multiple plant developmental stages. Our discovery of differentially abundant bacterial and fungal taxa in wild type finger millet and five cultivars provides further evidence for the microbiome-modulating effects of domestication. Further, we show that the microbial community structure of the root endosphere and rhizosphere changes as the plant advances through different growth stages. The unique bacterial and fungal species detected in each sample type and shared core microbiome provide further evidence of the selective pressure exerted during domestication. Moreover, the unique species found in the cultivars show that the endosphere microbiome is not merely drawn from the rhizosphere and some microbial species detected in the root compartments were not even present in the bulk soil, suggesting that they may have been passed down from the parent plant. This possibility warrants further investigation.

## Conclusion

An in-depth analysis of the microbial community structure in two root compartments – the rhizosphere and the endosphere – showed that the structures of their microbiomes change as the plant advances from one developmental stage to another. Moreover, the dynamics of the root-associated microbial community were shown to differ markedly between cultivars. Phylogenetic profiling revealed that the root bacteriota were dominated by the phyla *Proteobacteria*, *Actinobacteria*, and *Bacteroidetes* in both the seedling and flowering growth stages. Similarly, the fungal phyla *Ascomycota* and *Basidiomycota* were identified in all studied samples. It thus seems that these bacterial and fungal phyla comprise the core microbiomes in the studied environments. Despite this, their relative abundance varied between compartments, cultivars, and plant developmental stages. The endosphere was generally more diverse than the rhizosphere, especially in the flowering stage. The reduced diversity of the rhizosphere during the flowering stage was accompanied by pronounced enrichment of certain bacterial taxa, strongly suggesting that plants can select subsets of microbes in individual developmental stages, presumably for specific functional purposes. Moreover, plants can selectively enhance the enrichment of microbial genera from the core microbiota in individual compartments.

The differing community responses observed across plant developmental stages may reflect changes in the composition of the root exudates. These changes likely correspond to shifts in microbial community dynamics driven by the differing ability of various taxa to utilize specific substrates. Further study will be needed to test this hypothesis and determine the interactive functions of rhizocompartments and their microbiomes in different growth stages and plant varieties. Overall, this study has highlighted the shaping influence of both cultivar type and plant developmental stage on microbiome community profiles in root compartments.

## Acknowledgments

RRV acknowledge support from the Swedish Research Council for Environment, Agricultural Sciences and Spatial Planning (FORMAS; grant numbers 2019– 01316), Novo Nordisk Fonden (0074727), SLU Centre for Biological Control, the Swedish Research Council (2019–04270), SLU Partneskap Alnarp and Carl Tryggers Stiftelse för Vetenskaplig Forskning (CTS 20: 464).

## Availability of data and materials

All amplicon data used in this study were deposited into the sequence read archive (SRA) under BioProject accessionPRJNA1061570.

## Declarations

Not applicable.

## Ethics approval and consent to participate

Not applicable.

## Consent for publication

Not applicable.

## Supplementary Information

Additional file 1.

Relative abundance of each bacterial phylum from two rhizocompartments of six finger millet cultivars at two plant developmental stages.

Additional file 2.

Relative abundance of each bacterial genus from two rhizocompartments of six finger millet cultivars at two plant developmental stages.

Additional file 3.

Relative abundance of each fungal phylum from two rhizocompartments of six finger millet cultivars at two plant developmental stages.

Additional file 4.

Relative abundance of each fungal genus from two rhizocompartments of six finger millet cultivars at two plant developmental stages.

Additional file 5.

Core microbiome analysis of shared and unique bacterial and fungal species in the endosphere and rhizosphere of wild type plants (WT; Africana), five different cultivars (Wama, Tessema, Tadesse, Padet, and Axum), and control samples (soil without plantation) during the seedling and flowering growth stages.

Additional file 6.

Taxonomic data on the co-occurrence patterns of bacterial and fungal species in the endosphere and rhizosphere during the seedling and flowering growth stages.

